# Genome-wide association analysis identifies novel loci for chronotype in 100,420 individuals from the UKBiobank

**DOI:** 10.1101/038620

**Authors:** Jacqueline M. Lane, Irma Vlasac, Simon G. Anderson, Simon Kyle, William G. Dixon, David A. Bechtold, Shubhroz Gill, Max A. Little, Annemarie Luik, Andrew Loudon, Richard Emsley, Frank AJL. Scheer, Deborah A. Lawlor, Susan Redline, David W. Ray, Martin K. Rutter, Richa Saxena

## Abstract

Our sleep timing preference, or chronotype, is a manifestation of our internal biological clock. Variation in chronotype has been linked to sleep disorders, cognitive and physical performance, and chronic disease. Here, we perform a genome-wide association study of self-reported chronotype within the UKBiobank cohort (n=100,420). We identify 12 new genetic loci that implicate known components of the circadian clock machinery and point to previously unstudied genetic variants and candidate genes that might modulate core circadian rhythms or light-sensing pathways. Pathway analyses highlight central nervous and ocular systems and fear-response related processes. Genetic correlation analysis suggests chronotype shares underlying genetic pathways with schizophrenia, educational attainment and possibly BMI. Further, Mendelian randomization suggests that evening chronotype relates to higher educational attainment. These results not only expand our knowledge of the circadian system in humans, but also expose the influence of circadian characteristics over human health and life-history variables such as educational attainment.

---

Chronotype is a behavioral manifestation of our internal timing system, the circadian clock. Individual variation within our biological clock drives our morning or evening preferences, thereby making us into “morning larks” or “night owls”. Chronotype is influenced by many factors, including age, sex, social constraints, and environmental factors, among others^1^. Chronotype has been associated with sleep disorders, cognitive and physical performance, chronic metabolic and neurologic disease, cancer and premature aging,^2^ in particular when there is desynchrony between internal chronotype and external environment increasing disease risk^3^. Despite the importance of circadian rhythms to human health and their fundamental role demonstrated in model organisms,^4,5^ little is known about biological mechanisms underlying inter-individual variation in human chronotype or how it impacts on our health and physiology.

Genes that encode molecular components of the core circadian clock *(PER2, PER3)* or regulate the pace of the clock *(CSNK1D)* are disrupted in Advanced Sleep Phase Syndrome (ASPS) and Delayed Sleep Phase Syndrome (DSPS) both of which are monogenic circadian rhythm disorders causing extreme advance or delay in sleep onset^6^. ASPS mutations shorten circadian period in humans and mice^7,8^, linking the change in pace of the clock with sleep timing preference. More detailed biochemical and functional characterization of these mutations have greatly enhanced understanding mechanisms regulating the circadian clock. Emerging evidence suggests that subjects with ASPS may be at increased risk for chronic disease, such as cardio-metabolic disease^9^ or show familial segregation of the causal mutation with both advanced sleep phase and migraine^10^.

In addition to monogenic sleep phase disorders, pronounced inter-individual variation in chronotype exists within the general population^5^, and epidemiologic associations with adverse health outcomes have been reported^2,11^. Chronotype is heritable as estimated by twin and family studies (12-42%)^12-14^ but its genetic basis has not yet been well defined. Candidate gene association studies have reported variation associated with morningness or eveningness preference in the *CLOCK, PER1, PER2,* and *PER3* genes^15^; however, these studies have often had limited reproducibility, suffering from small sample sizes, heterogeneity in chronotype assessment and inadequate correction for population structure. Recently, a genome-wide association study (GWAS) for self-reported habitual bedtime identified variation in *NPSR1^12^,* but again robust replication of this finding has not been reported. Nonetheless, these studies suggest that novel genetic loci for chronotype, like for other complex traits, may be identified by GWAS provided that sufficiently large cohorts are used.

To define the spectrum of genetic variation contributing to variation in human circadian phenotype, and identify associative or causal links between chronotype and other health indices, we perform the largest genome-wide association study (GWAS) of self-reported chronotype to date, within the UK Biobank cohort (n=100,420), a unique resource with an extensive set of individual life history parameters. Self-reported chronotype has been validated in previous studies, and correlates significantly with objectively measured physiological rhythms^16^. Our work identifies several novel genetic loci that associate significantly with chronotype, and importantly reveals a significant genetic correlation between chronotype and schizophrenia risk, BMI, and educational attainment.

## Results

### Twelve genome-wide significant association signals

Variation in chronotype associated significantly with age, sex, sleep duration, depression and psychiatric medication use, with ‘eveningness’ being associated with younger age, being male, having a longer sleep duration, being more likely to be depressed or using psychiatric medication **(Supplementary Table 1).** These characteristics together explained 1.4% of variation in chronotype.

Two parallel primary GWAS analyses of genotyped and imputed SNPs were performed using regression models adjusting for age, sex, 10 principal components of ancestry and genotyping array: an ordinal score of chronotype based on 4 categories from “definite morning” to “definite evening” treated as a continuous trait, using the whole population (n=100,420) and a binary variable of chronotype extremes (8,724 definite evening type cases vs. 26,948 definite morning type controls), to enrich for rarer variants expected to have stronger effects. In total, 12 genome-wide significant loci were identified **(Figure 1-2, Table 1,** and **Supplementary Figure 1,** p<5 x 10^-8^) of which three surpassed genome-wide significance in both analyses **(Table 1).** Association was observed near *PER2,* an ASPS gene, and three other association signals were found in or near genes with a well-known role in circadian rhythms *(APH1A, RGS16,* and *FBXL13),* consistent with the hypothesis that circadian clock biology contributes to variation in chronotype. Conditional analyses at the 12 loci implicated one suggestive secondary association signal, a missense variant (V903I) in the core circadian clock gene *PER2* (p=8.43x10^-8^) predicted to be damaging (Polyphen 0.984, CADD scaled 16.21; **Supplementary Table 2);** thus, confirming that core circadian clock genes disrupted in ASPS harbor common variants that contribute to variation in chronotype. Together, in the discovery sample, the 12 loci explain 4.3% of variance in chronotype. Credible set analyses^17^ highlight a limited number of potential causal variants at each locus **(Table 1).**

**Table 1.** Genome-wide significant loci associated with chronotype in subjects of European ancestry in the UKBiobank.

|  |  |  |  |  | Continuous Chronotype (n=100,420)<br>(1-4 ranging from "definite morning"<br>to "definite evening") |  | Extreme Chronotype<br>(n= 8,724 evening type cases,<br>n=26,948 morning type controls) |  |  |
| --- | --- | --- | --- | --- | --- | --- | --- | --- | --- |
| SNP | Chr:position<br>NCBI 37 | Nearest<br>Gene | Alleles<br>(E/A) | EAf | Beta (SE) | p-val | OR (95% CI) | p-val | Most likely causal<br>SNPs (probability)* |
| rs141175086 | 1:780,397 | *LINC01128* | **C/T** | 0.998 | 0.266 (0.070) | $1.42 \times 10^{-4}$ | 2.16 [1.34-3.49] | **$4.38 \times 10^{-8}$** | rs141175086 (1.00) |
| rs2050122 | 1:19,989,205 | *HTR6* | **C/T** | 0.80 | 0.031 (0.006) | **$4.61 \times 10^{-8}$** | 1.12 [1.07-1.17] | $7.94 \times 10^{-7}$ | rs2050122 (0.385) |
| rs76681500 | 1:77,713,434 | *AK5* | **G/A** | 0.84 | 0.043 (0.006) | **$1.50 \times 10^{-12}$** | 1.16 [1.10-1.22] | **$1.77 \times 10^{-9}$** | rs76681500 (0.665) |
| rs10157197 | 1:150,250,636 | *APH1A* | **A/G** | 0.40 | 0.028 (0.005) | **$1.48 \times 10^{-9}$** | 1.13 [1.09-1.17] | **$1.27 \times 10^{-11}$** | rs10157197 (0.085),<br>rs11205355 (0.085) |
| rs1144566 | 1:182,569,626 | *RGS16* | **C/T** | 0.97 | 0.099 (0.013) | **$2.62 \times 10^{-14}$** | 1.35 [1.22-1.50] | **$1.29 \times 10^{-8}$** | rs1144566 (0.213),<br>rs694383 (0.213),<br>rs12743617 (0.213),<br>rs509476 (0.213) |
| rs11895698 | 2:239,338,495 | *ASB1* | **T/C** | 0.14 | 0.035 (0.006) | **$1.15 \times 10^{-8}$** | 1.10 [1.05-1.15] | $1.30 \times 10^{-4}$ | rs11895698 (0.279),<br>rs3769118 (0.279) |
| rs11708779 | 3:55,934,939 | *ERC2* | **G/A** | 0.65 | 0.026 (0.005) | **$3.08 \times 10^{-8}$** | 1.09 [1.06-1.13] | $1.20 \times 10^{-6}$ | rs11708779 (0.769) |
| rs148750727 | 4:188,022,952 | *FAT1* | **T/G** | 0.995 | 0.154 (0.033) | $3.61 \times 10^{-6}$ | 2.34 [1.69-3.23] | **$1.58 \times 10^{-8}$** | rs148750727 (1.00) |
| rs372229746 | 7:102,158,815 | *FBXL13* | **A/G** | 0.45 | 0.034 (0.005) | **$5.18 \times 10^{-10}$** | 1.12 [1.07-1.16] | $4.29 \times 10^{-7}$ | N/A |
| rs17311976 | 8:131,637,337 | *ADCY8* | **C/T** | 0.19 | 0.028 (0.006) | $1.08 \times 10^{-6}$ | 1.13 [1.08-1.18] | **$3.37 \times 10^{-8}$** | N/A |
| rs542675489 | 12:120,994,888 | *RNF10* | **C/CA** | 0.60 | 0.027 (0.005) | **$3.29 \times 10^{-9}$** | 1.09 [1.05-1.13] | $1.38 \times 10^{-5}$ | rs542675489 (1.00) |
| rs4821940 | 22:40,659,573 | *TNRC6B* | **C/T** | 0.55 | 0.026 (0.004) | **$1.05 \times 10^{-8}$** | 1.07 [1.04-1.11] | $8.56 \times 10^{-5}$ | rs4821940 (0.172) |
| Suggestive Secondary Signal |  |  |  |  |  |  |  |  |  |
| rs35333999 | 2:239161957 | *PER2* | **T/C** | 0.043 | 0.059 (0.011) | $8.43 \times 10^{-8}$ | 1.21 [1.11-1.32] | $9.01 \times 10^{-6}$ | N/A |

E=effect allele, A=alternative allele, Chr=chromosome, OR=Odds Ratio, CI=confidence interval. Ancestral allele is indicated in bold. EAF=effect allele frequency. Candidate Putative causal SNPs were identified by PICS. Note, increasing beta and Odds Ratio indicate greater evening type preference.

**Figure 1.**
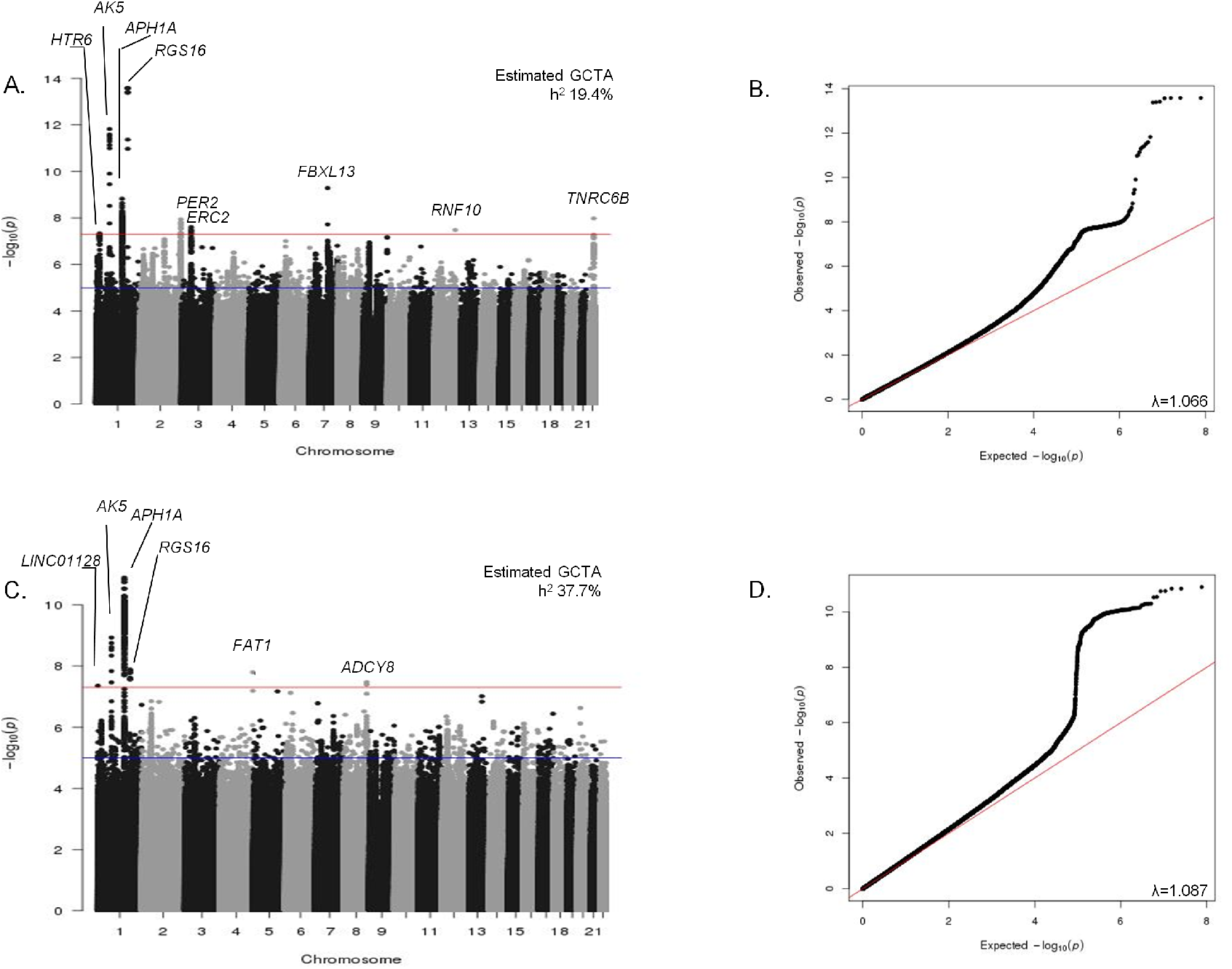
Manhattan and QQ plots for genome-wide association analysis of both continuous (A & B) and extreme (C & D) chronotype. Manhattan plots (A. and C.). Red line is genome-wide significant (5×10^-8^) and blue line is suggestive (1×10^-6^). Q-Q plots (B. and D.). Nearest gene name is annotated. Heritability estimates were calculated using BOLT-REML and lambda inflation values using GenABEL in R.

**Figure 2.**
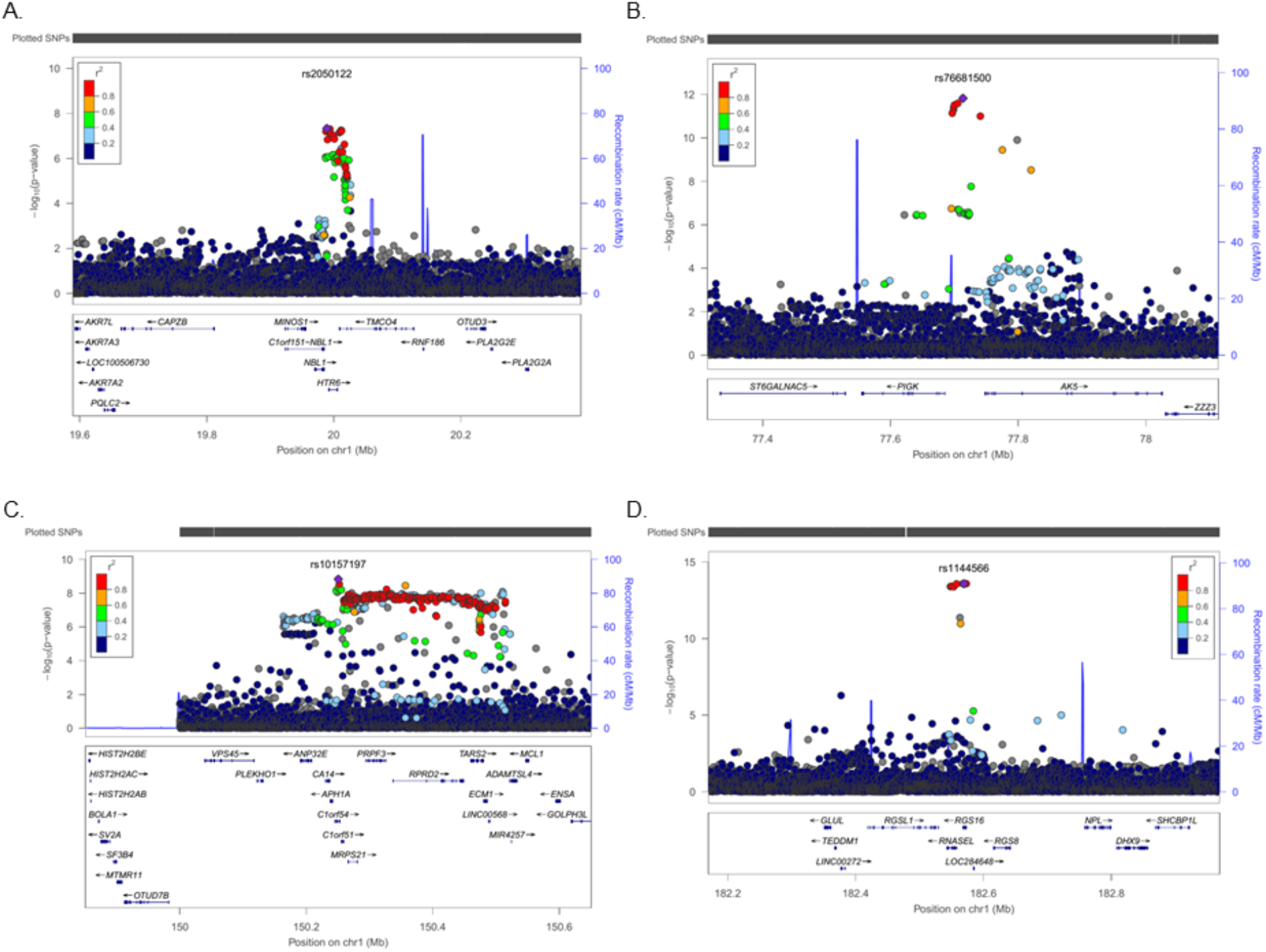
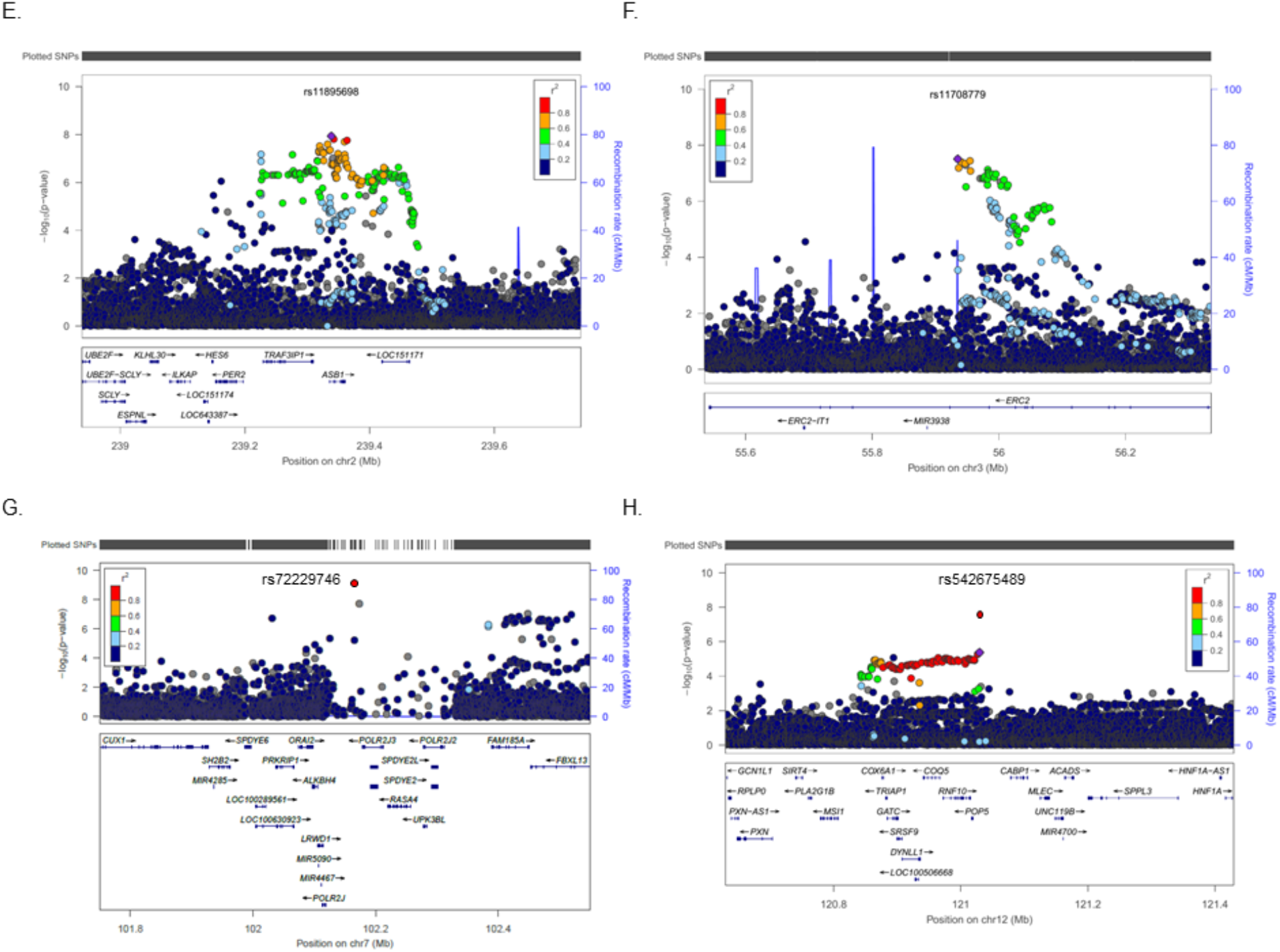
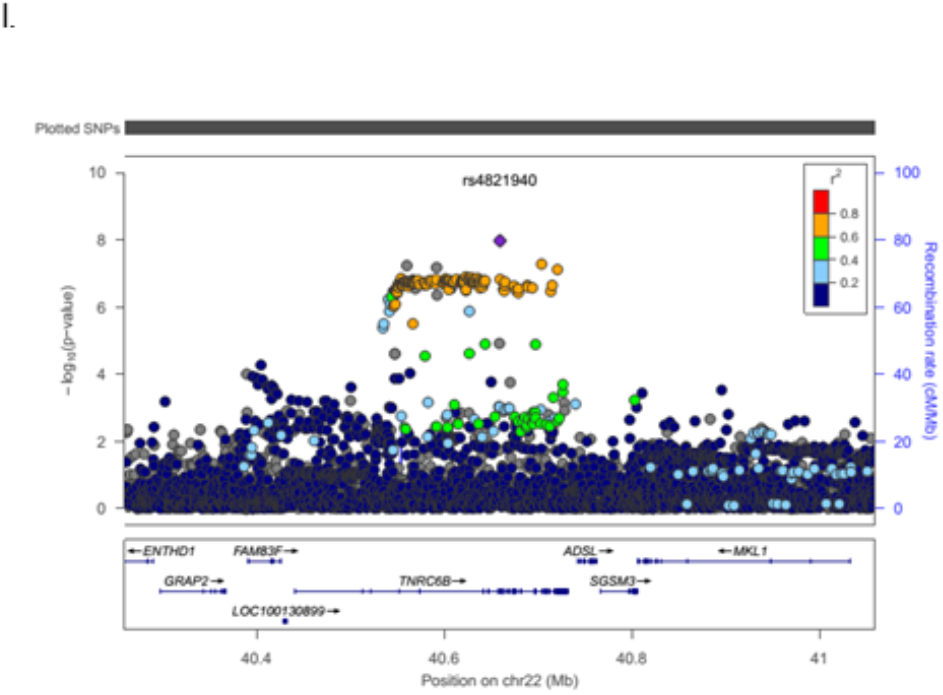
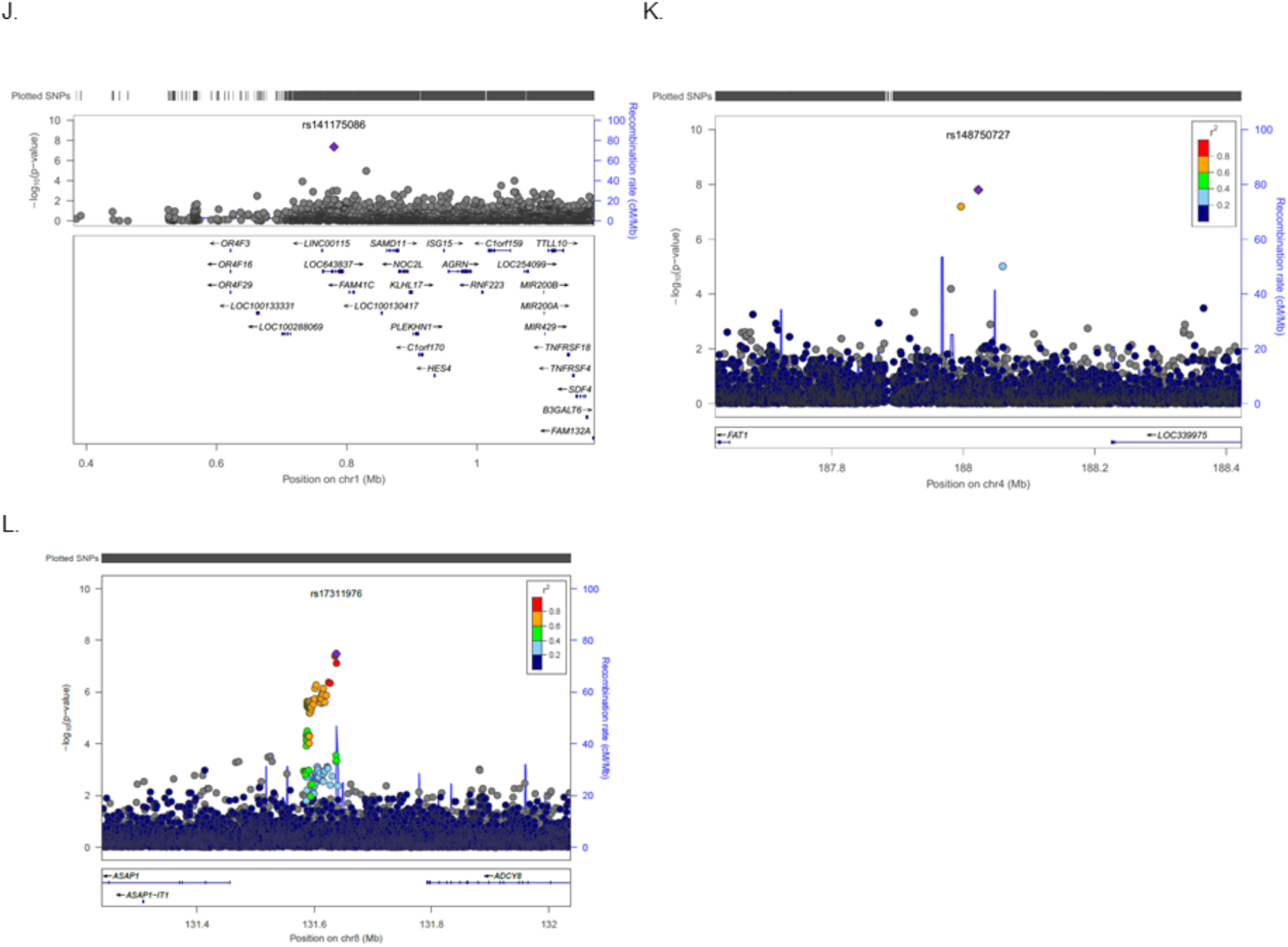
Regional association plots for genome-wide significant chronotype loci (A.-L.). Panels A-I show loci associated with continuous chronotype, J-L show loci associated with extreme chronotype. Genes within the region are shown in the lower panel. The blue line indicates the recombination rate. Filled circles show the -log10 P value for each SNP, with the lead SNP shown in purple. Additional SNPs in the locus are colored according to correlation (*r*^2^) with the lead SNP (estimated by LocusZoom based on the CEU HapMap haplotypes). *chr7 rs372229746 is not in the reference panel, therefore LD data is unavailable for this SNP.

Robustness of the self-reported chronotype trait and genetic loci identified here was further validated by an independent GWAS of extreme chronotype from Hu et al.^18^. 8 of 15 reported loci replicated in our study, and all 15 showed a consistent direction of effect in our study. Three additional loci attain genome-wide significance in meta-analysis of both studies using publicly available results for the 15 SNPs from Hu et al. (near genes *PER3, VIP* and *TOX3:* **Supplementary Table 3).**

No evidence of association was observed for previously reported SNPs from other candidate gene or GWA studies **(Supplementary Table 4).** The *PER3* VNTR (rs57875989) previously associated with chronotype^19^ was not directly genotyped or imputed in this study; nonetheless, a suggestive association signal was observed encompassing this region of *PER3* (lead SNP: rs7545893 p=6.5x10^-8^; 33KB from *PER3* VNTR and largely independent from the lead 23and me *PER3* region SNP (r^2^=0.186 in 1KG CE^18^U; **Supplementary Figure 2).**

Secondary analyses were performed on the 12 lead SNPs from chronotype loci, including 1) separate comparison of effects on morningness and eveningness, 2) sex-specific analysis, 3) pair-wise genetic interaction analysis, and 4) regression models including additional covariates. Comparison of case extremes (8,724 evening, or 26,948 morning) to the collapsed middle group (n= 64,748) revealed three loci *(LINC01128, APH1A, FAT1)* with stronger effects in the eveningness case-control analysis as opposed to morningness analysis. A rare variant (0.2%) at the *LINC01128* locus (rs141175086C) exhibited the most striking protective effect for eveningness (OR=0.22 (0.10-0.50), p=1.7x10^-5^) but only a small risk effect for morningness (OR=1.30 (0.97-1.75), p=0.08; **Supplementary Table 5).** No significant sex-specific effects **(Supplementary Table 6)** or epistasis between loci **(Supplementary Table 7)** were detected. Similarly, sensitivity analyses adjusting for factors known to associate with chronotype, including sleep duration and disorders, depression, and psychiatric medication use did not significantly alter the effect estimates or strength of the associations **(Supplementary Table 8).**

Candidate causal genes at these loci are highlighted in **Supplementary Note 1.** The 12 loci encompass 72 candidate genes enriched in pathways for circadian rhythms (p_adj_=0.014), mental disorders (p_adj_=0.001), sleep disorders (p_adj_=0.005), the spliceosome (p_adj_=0.020), and Alzheimer’s disease (p_adj_=0.030) among others **(Supplementary Table 9).** In addition, four loci are located in or near genes with a well-known role in circadian rhythms *(PER2, APH1A, RGS16,* and *FBXL13),* however whether these genes are responsible for the association signals observed remains to be established. The remaining eight loci offer the potential of novel biological insights into circadian rhythms **(Supplementary Note 1).** Several candidate causal genes have been implicated in circadian rhythms. *TNRC6B* controls circadian behavior in flies^20^ and is bound by known circadian transcription factors. *MCL1* has rhythmically expressed mRNA in liver^21^, disrupts circadian rhythms in an RNAi screen using a human osteosarcoma cell line ^22^, and is bound by known circadian transcription factors ^23^. *HTR6 is* a G-protein coupled receptor known to regulate the sleep wake cycle^24-26^.

Fine-mapping, sequencing and experimental studies are necessary to identify the causal gene(s) and variant(s) at each locus in order to understand mechanisms by which DNA variation influences variation in chronotype. However, clues may emerge from exploration of bioinformatic annotations of candidate regulatory variants and ENCODe analyses of chromatin states and bound proteins^27^. For example, rare variant rs141175086 is predicted to disrupt a binding site for the known circadian transcription factor DEC1 in an enhancer element within or upstream of previously uncharacterized lincRNAs *(LOC643837, LINC01128).*

### Pathway analyses

Heritability of chronotype, captured by genome-wide genotypes in this study, was estimated to be 19.4% (continuous) and 37.7% (extreme) using GCTA^28^. Heritability partitioning of continuous chronotype GWAS by tissue and functional category using LD-score regression^29^ identified enrichment in the central nervous system (Enrichment 2.63, p=1.91x10^-6^) and adrenal/pancreatic tissues (Enrichment 3.63, p=1.34x10^-8^; **Figure 3a** and **Supplementary Table 10).** Regions of the genome annotated as highly conserved across mammals^30^ (Enrichment 14.33, *p*=1.75x10^-9^), and in regions of histone 3 lysine 4 monomethylation that mark active/poised enhancer elements (Enrichment 1.30, p=0.0017; **Figure 3a** and **Supplementary Table 10)** were significantly enriched, supporting a key role of circadian rhythms throughout mammalian evolution.

**Figure 3.**
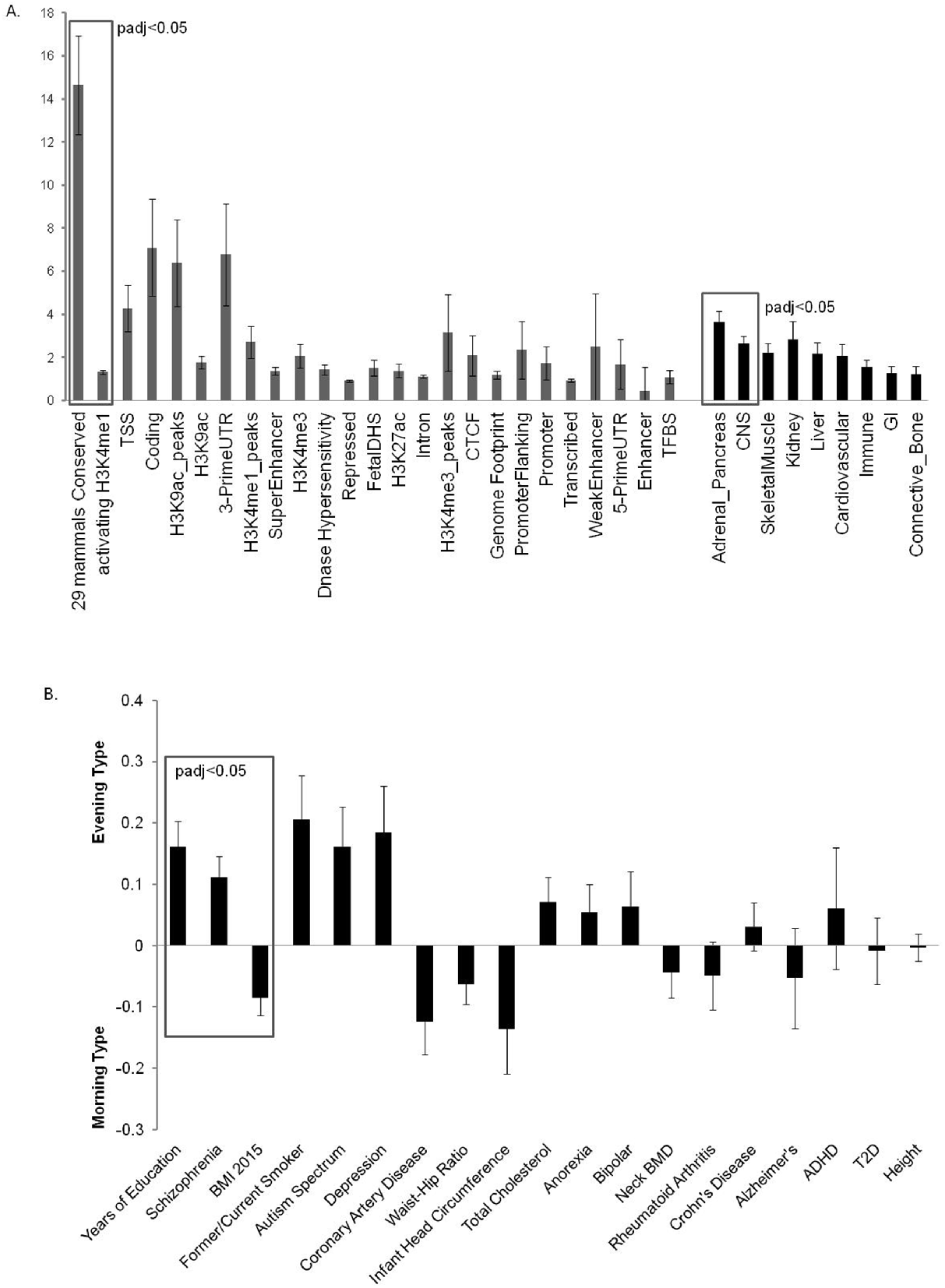
Overall genetic architecture of chronotype across tissues, functional categories, and cross-trait genetic correlation. A. Enrichment estimates for the main annotations and tissues of LDSC. Error bars represent 95% confidence intervals around the estimate. Categories are sorted by p-value, with boxes indicating annotations or tissues which pass the multiple testing significance threshold. B. Chronotype regression estimates of genetic correlation with the summary statistics from 19 publicly available genome-wide association studies of psychiatric and metabolic disorders, immune diseases, and other traits of natural variation. The horizontal axis indicates the phenotype compared to categorical chronotype and the vertical axis indicates genetic correlation. Error bars are standard errors. Abbreviations: Tss=transcription start site, DHS=DNase hypersenstivity, UTR=untranslated region, CTCF=CCCTC-binding transcription factor, TFBS=transcription factor binding site, CNS=central nervous system, GI=gastrointestinal, BMI=body mass index, ADHD=attention deficit hyperactivity disorder, T2D=type 2 diabetes.

Gene-based analysis^31^ identified 23 genes significantly associated with chronotype (p<2.8x10^-6^, **Supplementary Table 11, Supplementary Figure 3).** Pathway analysis^32^ shows a significant enrichment in this gene-set for genes previously implicated in Alzheimer’s disease (p_adj_=0.0176) and dementia (p_adj_=0.0192), eye abnormalities (p_adj_=0.0176) and eye diseases (p_adj_=0.0253), chromosomal deletions (p_adj_=0.0253), and brain diseases (p_adj_=0.0253), central nervous system diseases (p_adj_ =0.0253), and mental disorders (p_adj_=0.0365). In support, integrative analysis of signals with p<1 x 10^-5^ using DEPICT^33^, a tool that uses predicted gene functions to prioritize genes, gene-sets and tissues, showed suggestive enrichment in gene-sets associated with ‘fear-response’ and ‘behavioral defense response’ (FDR<0.20), and central nervous and Hemic/ immune system tissues **(Supplementary Table 12).** In total, pathway analyses link the genetics of chronotype to central nervous system function and neurological disorders including dementia and affective disorders.

### Genetic links with schizophrenia and educational attainment

Given that circadian rhythms play a fundamental role in human physiology, a key question is the extent to which the genetics of chronotype is shared with other behavioral or disease states, and importantly whether genetic relationships between chronotype and other traits are causal. To address this, we tested for genetic correlation of chronotype with GWAS variants for 19 phenotypes spanning a range of cognitive, neuro-psychiatric, anthropometric, cardio-metabolic and auto-immune traits using LD score regression on chronotype GWAS and publicly available GWAS for each trait^34^. Genetic correlations suggested that tendency towards an evening chronotype is related to greater years of education (r_g_ (SE) 0.161 (0.041), p=8.96 x 10^-5^) and increased schizophrenia risk (r_g_ (SE) 0.112 (0.034), p=0.0011 **(Figure 3b** and **Supplementary Table 10).** Genetic correlations also suggested that a morning chronotype may share underlying biology with increased BMI (r_g_ (SE) −0.0851 (0.0281), p=0.0025; **Figure 3b** and **Supplementary Table 10).**

### Mendelian randomization analyses

To explore whether the relationship between chronotype and traits with significant genetic correlations might be causal, we tested for association of a risk score of genome-wide significant chronotype SNPs from 23andMe^18^ with years of education, schizophrenia and BMI. SNPs can be used as instrument variables to test for a causal relationship between two traits, and because genotypes are assigned randomly at meiosis, genetic association is not biased by confounding or reverse causation possible in observational epidemiology^35,36^. Since individuals do not know their genotype any phenotypic misclassification will be random with respect to genotype. In UKBiobank, a significant association was observed between a chronotype genetic risk score of SNPs related to eveningness and increased educational attainment (p=0.0167), but not schizophrenia (p=0.101) or BMI (p=0.285; **Supplementary Table 13).** Further instrumental variable analyses suggested that for each increase in ‘eveningness’ category, educational attainment increased by 7.5 months *(p*=0.021) **(Figure 4, Supplementary Table 13).** We then tested for reverse causation by assessing whether variation in education, schizophrenia, or BMI might cause variation in chronotype by testing for association of risk scores for each of these traits obtained from prior large-scale GWAS studies with chronotype. No significant associations were observed **(Supplementary Table 13).**

**Figure 4.**
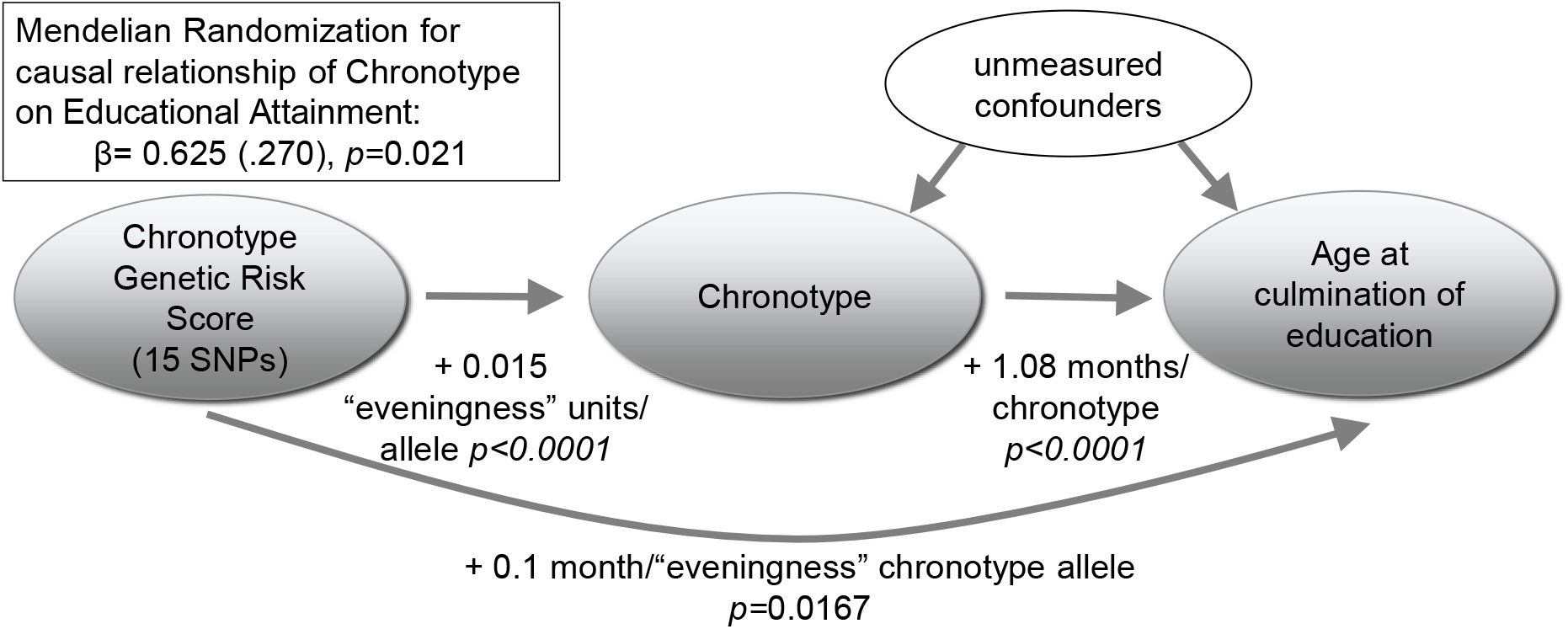
Mendelian Randomization: Under the assumptions of instrumental variable Mendelian randomization analyses^70^, our results show that having an evening chronotype results in higher educational attainment. In this analysis, for the chronotype risk score (comprised of 15 SNPs from the 23andMe GWAS of chronotype, weighted by effect size), the β coefficient for the association with chronotype was regressed on the β coefficient for the association with the main educational attainment trait in the UKBiobank (n=68,718) using TSLS.

## Discussion

In this largest GWAS of chronotype to date, we report the discovery of twelve genetic loci associated with chronotype, and pathway analysis suggests key roles of genes in the nervous and ocular systems. Further, we demonstrate shared biology of chronotype with schizophrenia, and possibly BMI, with a putative causal link to educational attainment.

Several lines of evidence support these association signals as true positives that may help to uncover new aspects of circadian biology in humans. First, we detect signals in or near known circadian genes at 4 of the 12 loci, including near and in *PER2,* a clock gene previously associated with ASPS^6^. Second, 3 of these signals have been observed in an independent GWAS^18^ suggesting independent validation of our findings. Third, novel associated loci include candidate central circadian clock genes with rhythmic expression in the SCN or circadian behavioral phenotypes in model organisms. Fourth, genes under association peaks are enriched for central nervous system and ocular processes, both important for generation of circadian rhythms. Additional replication to confirm chronotype genetic associations and functional follow-up will be necessary to identify causal genes and circuits disrupted by causal variants at these loci.

Our study also defines the genetic architecture of self-reported chronotype, revealing heritability estimates consistent with previous literature^12-14^, despite using a different questionnaire instrument than previous studies^16^. The 12 genome-wide significant loci appear to explain a large fraction of chronotype variance (4.3%) but this may be over-estimated due to winners curse, or may reflect lower polygenicity of chronotype than seen for other complex traits, since variation in a limited number of biological processes (light-sensing, core circadian clock and limited downstream effectors) may be causal. Significant enrichment of heritability in highly conserved regions is consistent with the strong conservation of circadian rhythms throughout evolution^37^ and may aid in fine-mapping of causal variants and creation of faithful animal models for future experimental studies. Similarly, enrichment of heritability in activating enhancer sites and borderline enrichment in transcriptional start sites is consistent with the role of the circadian molecular clock in fine-tuning of transcriptional regulation^23^.

The association signals at loci identified by our study when combined with signals from 23andme cover genes identified in GWAS for restless legs syndrome and Mendelian and model organism studies of narcolepsy, suggesting overlap with other sleep traits. Genetic variants in the region of *TOX3* have been previously associated with restless legs syndrome (RLS) in a GWAS^38^. Although the chronotype-associated variant (rs12927162) is not in linkage disequilibrium with the lead RLS variant (rs3104767; r^2^<0.001 in 1KG CEU population), it does suggest that TOX3 may have a broader role in basic sleep/circadian physiology. Additionally, rare forms of severe early onset narcolepsy in humans^39^ and familial narcolepsy in canines^40^ re caused by mutations in *HCRTR2,* again suggesting shared underlying biology.

Chronotype has previously been associated with many behaviors and diseases, such as cardiovascular disease, type 2 diabetes, metabolic disorders, risk-taking behavior, cancer, psychiatric disorders, and even creativity^1,3^. Comparing the genetic architecture of chronotype captured in this study with an initial series of select phenotypes with publicly-available GWAS data, we identified significant genetic overlap between chronotype and schizophrenia, educational attainment and possibly BMI. Previous literature links evening chronotype with schizophrenia^41-43^, consistent with our findings. These studies also demonstrate severe circadian sleep/wake disruptions in people with schizophrenia, indicating that this relationship may be bidirectional. However, our Mendelian randomization analyses did not support causal relationships between these two. It is possible that even with our large sample size, we are underpowered to rule out an effect of schizophrenia on chronotype.

We detect a surprising putative genetic link between morning chronotype and higher BMI. Previous observational studies have shown association of evening chronotype with higher BMI, poorer dietary habits, and decreased inhibitions^44-47^. Consistently, we noted an observational association between eveningness chronotype and BMI (beta=1.003 BMI units/chronotype; p=1x10^-4^; r= 0.011). Our genetic correlation analyses suggest the intriguing possibility that some underlying pathways contributing to morning chronotype might increase BMI. We acknowledge that independent replication and further large studies are required to fully understand the relationship between chronotype and BMI.

Until now, it has been difficult to discern causal relationships between chronotype and other traits because of the potential bias due to confounding or reverse causality, which are unlikely to affect genetic studies^48^. Our work suggests that tendency to eveningness chronotype is potentially causally related to increased educational attainment, but replication of these findings, and more comprehensive assessment of potential sources of bias will require future investigation. Previous studies have reported that night owls earn a larger mean income than their earlier rising counterparts^49^. Another study, performed at a top-ranked business school, demonstrated higher GMAT scores in evening types even within a high achieving group^50^. It is possible that there is misclassification in our self-reported measurement of chronotype. Whilst the question clearly asks for preference, participants might have been influenced by the reality of their working lives. Those from more deprived socioeconomic positions might have occupations that are more restrictive in terms of working hours and hence less able to ‘adhere’ to their preference. If this results in a relationship between socioeconomic position and misclassification then socioeconomic position would confound any observational associations. However, since participants are extremely unlikely to know their genotype for the variants we have identified, any misclassification of chronotype by genotype will be random with the expectation that the genetic correlation and Mendelian randomization studies would be biased towards the null.

Our study is well-powered to detect genetic variants associated with chronotype, with previous studies demonstrating the power of a sample size >100,000 for detecting genetic effects^51^. The study uses a single harmonized question across a large cohort, which is in contrast with previous studies that needed to harmonize data across several cohorts with varying measures of chronotype. Our measure of chronotype is based on self-identification, and may reflect timing preference more so than objective measures of chronotype and since it does not take weekday and weekend behavior into account, any misclassification may be related to occupation and/or socioeconomic position. However, as noted above, for our genetic correlation and Mendelian randomization analyses this would be expected to bias findings towards the null. Our cohort is aged 40 to 69 and of European ancestry, which reduces the likelihood of bias due to population structure, but means we cannot necessarily assume our results generalize to other groups. That said the distribution of chronotype is consistent with that found in previous studies^52-54^.

In summary, in a large-scale GWAS of chronotype, we identified 12 new genetic loci that implicate known components of the circadian clock machinery and point to previously unstudied genetic variants and candidate genes that might modulate core circadian rhythms or light-sensing pathways. Furthermore, genome-wide analysis suggests that chronotype shares underlying genetic pathways with educational attainment, schizophrenia and possibly BMI, and that evening chronotype might be causally related to higher educational attainment. This work should advance biological understanding of the molecular processes underlying circadian rhythms, and open avenues for future research in the potential of modulating circadian biology to aide prevention and treatment of associated diseases.

## Methods

### Population and study design

Study participants were from the UK Biobank study, described in detail elsewhere^55^. In brief, the UK Biobank is a prospective study of >500,000 people living in the United Kingdom. All people in the National Health Service registry who were aged 40-69 and living <25 miles from a study center were invited to participate between 2006-2010. In total 503,325 participants were recruited from over 9.2 million mailed invitations. Self-reported baseline data was collected by questionnaire and anthropometric assessments were performed. For the current analysis, individuals of non-white ethnicity were excluded to avoid confounding effects.

### Chronotype and covariate measures

Study subjects self-reported chronotype, sleep duration, depression, medication use, age, and sex on a touch-screen questionnaire. Chronotype was derived from responses to a chronotype question that participants answered, along with other study questions, on a touch-screen computer at each assessment centre. The question was taken from the Morningness-Eveningness questionnaire; it is the question from that questionnaire that explains the highest fraction of variance in preferences in sleep-wake timing and is an accepted measure of chronotype^54^. The question asks: “Do you consider yourself to be…” with response options “Definitely a ‘morning’ person”, “More a ‘morning’ than ‘evening person”, “More an ‘evening’ than a ‘morning’ person”, “Definitely an ‘evening’ person”, “Do not know”, “Prefer not to answer”. This question specifically does not ask about actual sleeping pattern, and nor does it distinguish between weekday and weekend behavior and was accessed at the time of exam, which crosses days of the week and seasons across participants. 498,450 subjects answered this question, but only the 153,000 with genetic data were considered for this analysis. Subjects who responded “Do not know” or “Prefer not to answer” were set to missing. Chronotype was treated both as a continuous trait, with chronotype coded 1-4, where 1 represents definite morning chronotype, and a dichotomous trait, with definite morning responders set to control (n=125,052) and definite evening responders set to case (n=41,741). Depression was reported in answer to the question “How often did you feel down, depressed or hopeless mood in last 2 weeks?” (cases, n=4,279). Subjects with self-reported shift work (n=22,165) or sleep medication use (n=4,575) were excluded.

### Genotyping and quality control

Of the ∽500,000 subjects with phenotype data in the UK Biobank, ∽153,000 are currently genotyped. Genotyping was performed by the UK Biobank, and genotyping, quality control, and imputation procedures are described in detail here^56^. In brief, blood, saliva, and urine was collected from participants, and DNA was extracted from the buffy coat samples. Participant DNA was genotyped on two arrays, UK BiLEVE and UKB Axiom with >95% common content and genotypes for ∽800,000 SNPs were imputed to the UK10K reference panel. Genotypes were called using Affymetrix Power Tools software. Sample and SNP quality control were performed. Samples were removed for high missingness or heterozygosity (480 samples), short runs of homozygosity (8 samples), related individuals (1,856 samples), and sex mismatches (191 samples). Genotypes for 152,736 samples passed sample QC (∽99.9% of total samples). SNPs were excluded if they did not pass QC filters across all 33 genotyping batches, with a missingness threshold of 0.90. Batch effects were identified through frequency and Hardy-Weinberg equilibrium tests (p-value <10^-12^). Before imputation, 806,466 SNPs pass QC in at least one batch (>99% of the array content). Population structure was captured by principal component analysis on the samples using a subset of high quality (missingness <1.5%), high frequency SNPs (>2.5%) (∽100,000 SNPs) and identified the sub-sample of European descent. Imputation of autosomal SNPs was performed to a merged reference panel of the Phase 3 1000 Genome Project and the UK10K using IMPUTE3^57^. Data was prephased using SHAPEIT3^58^. In total, 73,355,677 SNPs, short indels and large structural variants were imputed. Post-imputation QC was performed as previously outlined and an info score cut-off of 0.1 was applied. For GWAS, we further excluded SNPs with MAF <0.00016, a threshold which represents a minimum 50 counts of each genotype, a conservative threshold. In total, 100,400 samples of European descent with high quality genotyping and complete phenotype/covariate data were used for these analyses. Genotyping quality of two significant rare SNPs (rs1144566 and rs35333999) was verified by examination of genotyping intensity cluster plots **(Supplementary Figure 5).** In addition, for two significant imputed rare SNPs which checked Information Quality Scores (info) and found these to be above the standard threshold of 0.40 used to indicate good imputation quality^59^ (rs141175086 info =0.48 and rs148750727 info = 0.88). Considering the size of the genotyped UK Biobank cohort (N∽150,000), an information measure of 0.4 on a sample of 150,000 individuals indicates that the amount of data at the imputed SNP is roughly equivalent to perfectly observed genotype data in a sample of N∽60,000.

### Statistical Analysis

Genetic association analysis was performed in SNPTEST^60^ with the “expected” method using an additive genetic model adjusted for age, sex, 10 principal components of ancestry and genotyping array. Genome-wide association analysis was performed separately for continuous chronotype and “extreme” chronotype with a genome-wide significance threshold of 5x10^-8^. Follow-up analyses on genome-wide significant loci included sex interaction testing using a linear regression model including a sex*SNP interaction term, performed in R^61^, conditional analysis using SNPTEST conditioning on the lead signal in each locus ±500kb, covariate sensitivity analysis individually adjusting for sleep duration, sleep disorders, insomnia, and depression/psychiatric medication use. Heritability was calculated using BOLT-Reml^62^. Post-GWAS analysis of LD Score Regression (LDSC)^29,34,63^ was conducted using all UK Biobank SNPs also found in HapMap3^64^ and included publicly available data from 19 published genome-wide association studies, with a p-value threshold of 0.0026 after Bonferroni correction for all 19 tests performed. Gene-based testing was performed using VEGAS^31^ on GWAS summary statistics from SNPs and samples passing rigorous quality control, and gene-set enrichment of genes significant after Bonferroni correction was performed using Web-Gestalt^32^. Given that gene-based tests like VEGAS are sensitive to missing data and may show inflation and low power with if data for rare variants is missing^65^, we note low missingness rates (>80% of SNPs had over 99.5% genotyping call rate), and for <5% of SNPs that may have failed in a subset of 33 batches, imputation was used to infer missing genotypes. Furthermore, our gene-based testing included only single SNP association results for variants with over 50 minor allele counts. A Q-Q plot of inflation-adjusted gene-based results for 17,791 genes is shown in **Supplementary Figure 3.** Pathway-based analysis to identify enrichment in biological processes, gene-sets and tissues suggested by the top loci was performed in DEPICT^33^ for all SNPs present in 1KG phase 3^66^.

For Mendelian randomization analyses, the weighted genetic risk score was calculated by summing the products of the chronotype risk allele count for 15 SNPs multiplied by the scaled chronotype effect reported by 23andMe^18^ i.e. using weights from an independent study to our own). The instrumental variable analyses were performed in R^40^ using the two-stage-least-squares method (TSLS function in the SEM package). The risk scores for education, schizophrenia, and BMI were constructed using the GWS SNPs and weights from previously published GWAS^67-69^ and tested on chronotype using the summary statistics from our reported GWAS using the GTX package in R.

## Author Contributions

The study was designed by JML, MKR, and RS. JML, IV and RS performed genetic analyses. JML and RS wrote the manuscript and all co-authors helped interpret data, reviewed and edited the manuscript, before approving its submission. RS is the guarantor of this work and, as such, had full access to all the data in the study and takes responsibility for the integrity of the data and the accuracy of the data analysis.

## Acknowledgements

This work was supported by NIH grants R21HL121728-02 (RS), F32DK102323-01A1 (JML), R01HL113338-04 (JML, SR, and RS), The University of Manchester (Regional Innovation Funding) and UK Medical Research Council MC_UU_12013/5 (DAL). We would like to thank the participants and researchers from the UKBiobank who contributed or collected data. Data on glycemic traits have been contributed by MAGIC investigators and have been downloaded from www.magicinvestigators.org. Data on coronary artery disease / myocardial infarction have been contributed by CARDIo-GRAMplusC4D investigators and have been downloaded from www.CARDIOGRAMPLUSC4D.ORG. We thank the International Genomics of Alzheimer’s Project (IGAP) for providing summary results data for these analyses.

The authors have no competing financial interests to declare.

